## Supplementary Information for "Endosomal dysfunction contributes to cerebellar deficits in spinocerebellar ataxia type 6"

Anna A. Cook[1], Tsz Chui Sophia Leung[1], Max Rice[1,2], Maya Nachman[1], Élyse Zadigue-Dubé[1], Alanna J. Watt[1*]

[1]Biology Department, McGill University, Montreal, QC, Canada.

[2]Department of Biological Sciences, Columbia University, New York, NY, USA.

### Supplemental figures

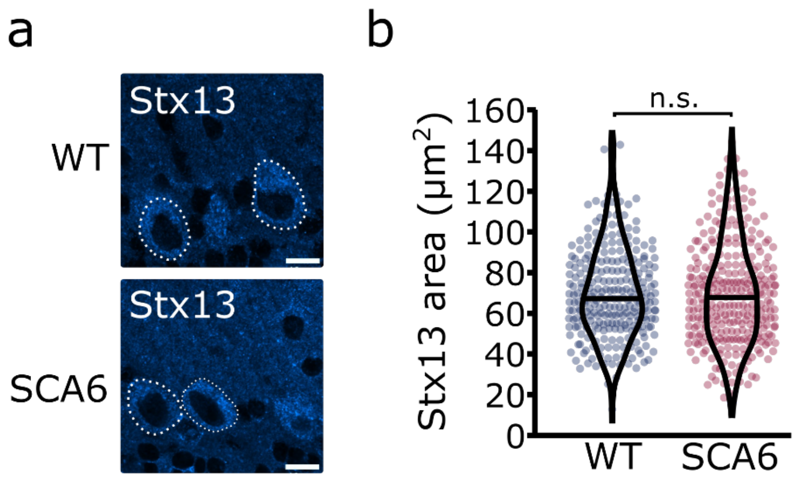


**Supplementary figure 1: A second Stx13 antibody confirms that recycling endosome area is unchanged in SCA6 Purkinje cells.**

(a) Representative image of the recycling endosome marker Stx13 within Purkinje cells of the anterior vermis. Scale bar, 10µm. (b) The area covered by Stx13 staining in Purkinje cells is unchanged between WT and SCA6 mice ($P = 0.59$; Mann Whitney *U* test; N = 4 WT mice, 268 cells; 6 SCA6 mice, 304 cells). n.s. $P > 0.05$.

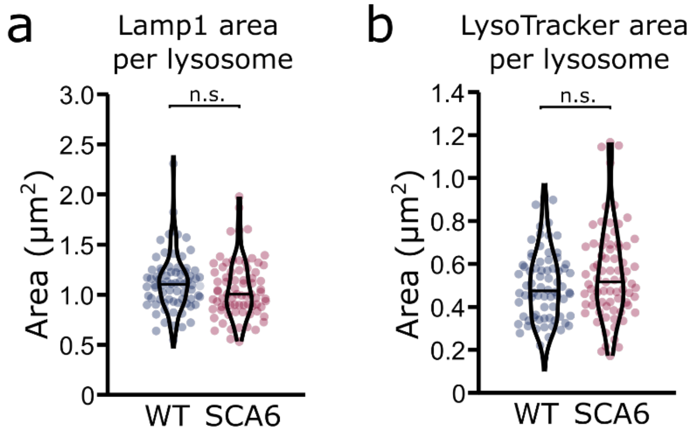


**Supplementary figure 2: Lysosome size is unchanged in SCA6 Purkinje cells.**

(a) The area covered by Lamp1 staining divided by the number of Lamp1 positive organelles is not significantly different in the Purkinje cells of WT and SCA6 mice ($P = 0.093$; N = 3 WT mice, 78 cells; 3 SCA6 mice, 76 cells). (b) The area covered by LysoTracker staining divided by the number of LysoTracker positive organelles is not significantly different in the Purkinje cells of WT and SCA6 mice ($P = 0.14$; N = 3 WT mice, 3 SCA6 mice). Mann Whitney *U* test used for all statistical comparisons; n.s. $P > 0.05$.

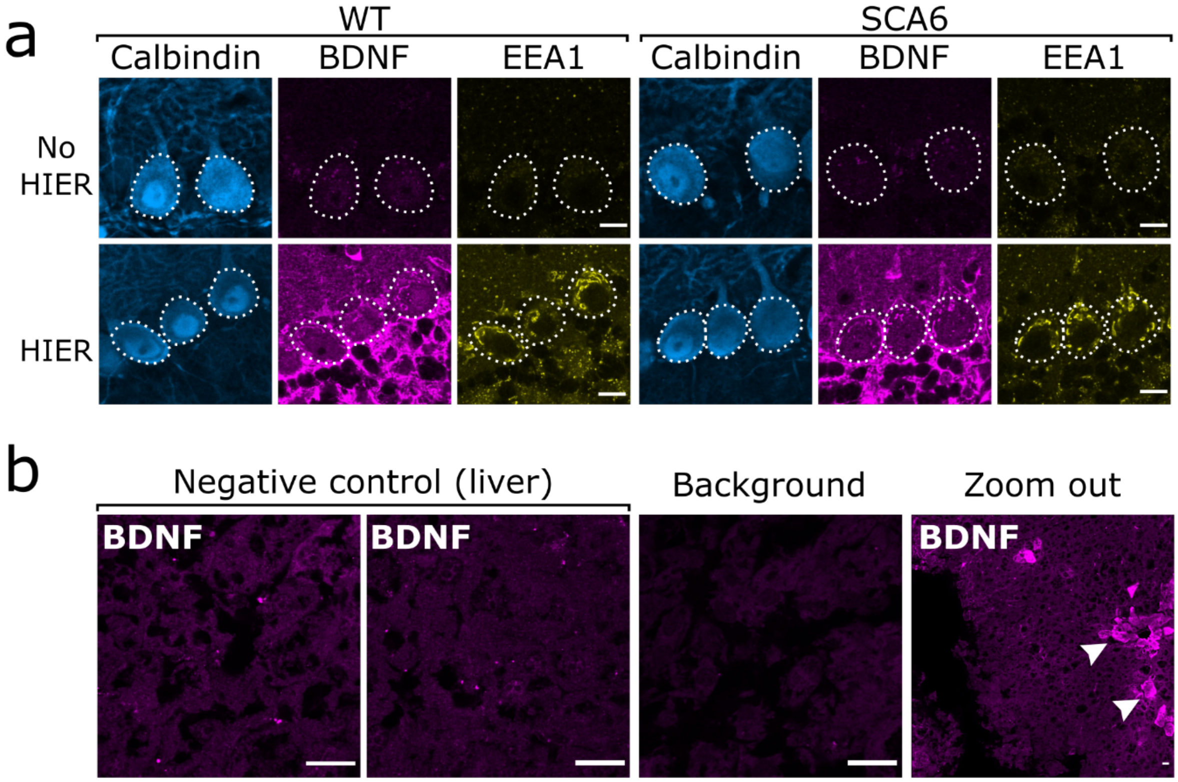


**Supplementary figure 3: BDNF immunostaining in the cerebellum requires heat-induced epitope retrieval (HIER) and is not detectable in the majority of liver tissue.** (a) HIER greatly enhances the signal from both BDNF and EEA1 staining in cerebellar vermis tissue from WT and SCA6 mice. Calbindin staining is unaffected. Scale bars, 10µm. (b) BDNF staining is undetectable in the majority of liver tissue from WT mouse, with only the BDNF-positive putative cholangiocytes (arrowheads) showing immunoreactivity with the BDNF antibody. Background slice was incubated without primary antibody but all other staining steps proceeded as normal. Scale bars, 20µm.

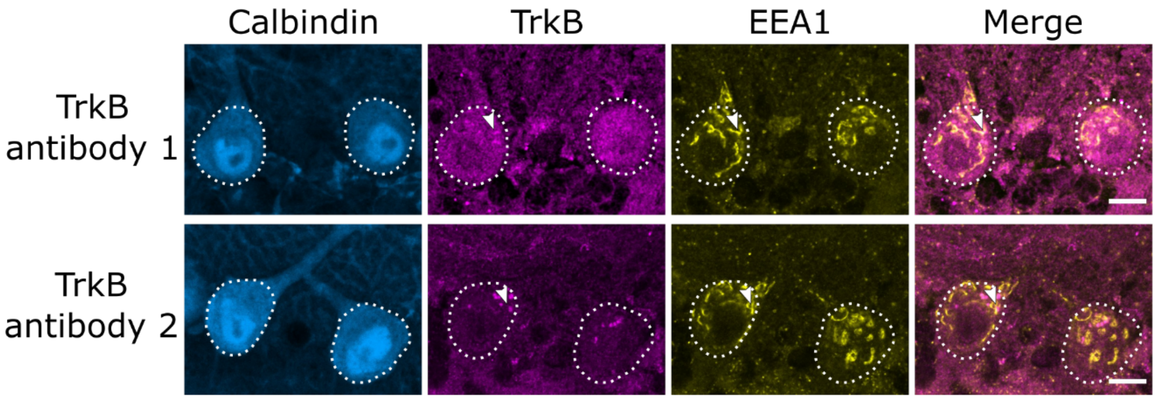

**Supplementary figure 4: A second TrkB antibody confirms endosomal localization of TrkB.** (a) Comparison of cerebellar vermis slices from the same WT mouse stained in the same batch show similar patterns of staining with two different TrkB antibodies. Arrowheads indicate TrkB colocalization with EEA1. Scale bars, 10μm.

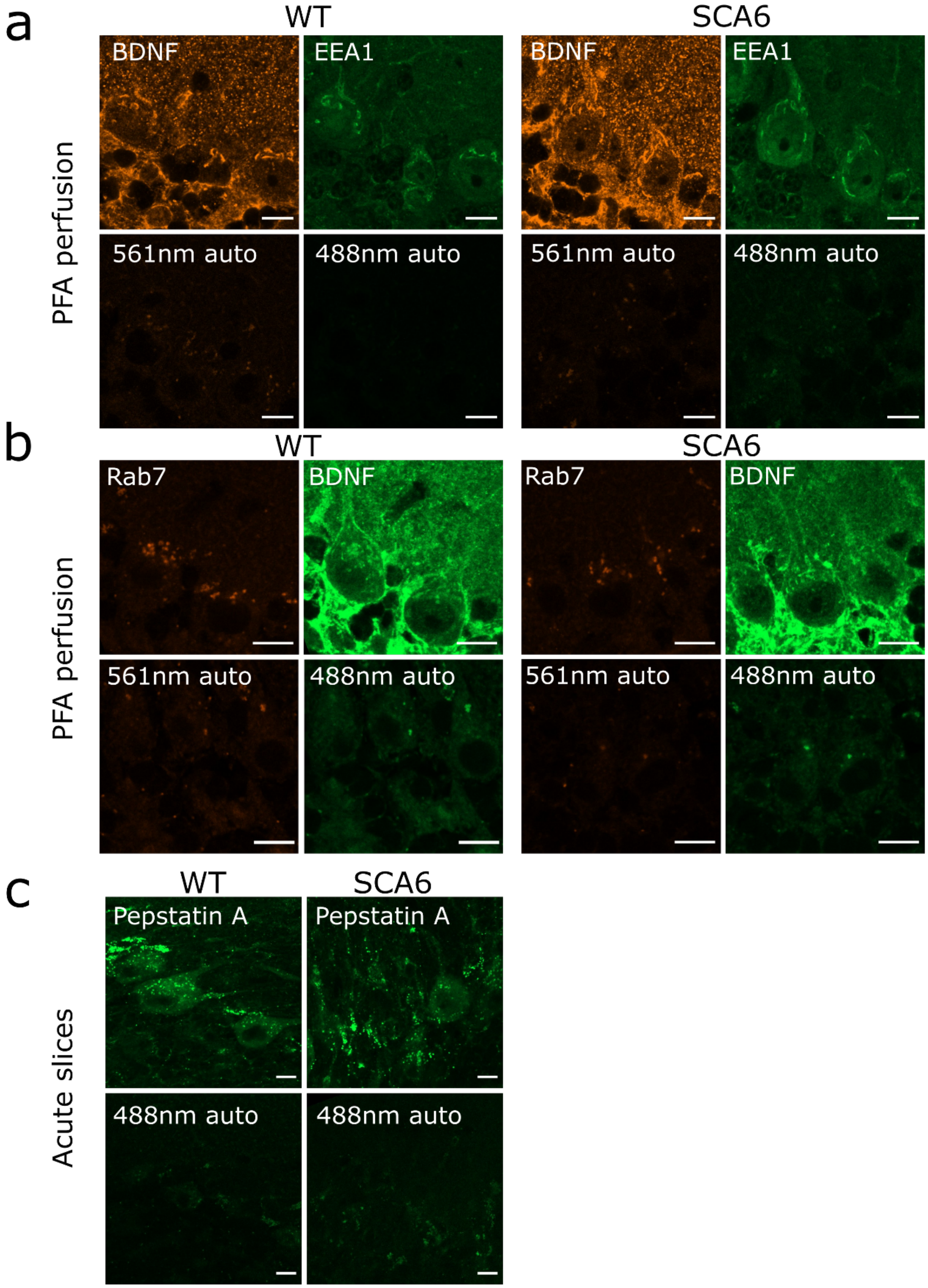

**Supplementary figure 5: Autofluorescence granules in Purkinje cells do not interfere with endosome identification.**

(a) Top row: raw images of BDNF and EEA1 staining in anterior vermis. Bottom row: raw images of slices from the same mice with no primary antibody incubation, showing autofluorescence from excitation at 561nm and 488nm. (b) Top row: raw images of Rab7 and BDNF staining in anterior vermis. Bottom row: raw images of slices from the same mice with no primary antibody incubation, showing autofluorescence from excitation at 561nm and 488nm. (c) Top row: raw images of Pepstatin A BODIPY FL incubation on acute slices that were post-fixed in PFA. Bottom row: raw images of slices from the same mice with no Pepstatin A incubation, showing autofluorescence from excitation at 488nm. Scale bars, 10µm.
